# Host immunity increases *Mycobacterium tuberculosis* reliance on cytochrome *bd* oxidase

**DOI:** 10.1101/2020.08.21.260737

**Authors:** Yi Cai, Eleni Jaecklein, Jared Mackenzie, Kadamba Papavinasasundaram, Andrew J. Olive, Xinchun Chen, Adrie Steyn, Christopher Sassetti

## Abstract

In order to sustain a persistent infection, *Mycobacterium tuberculosis* (*Mtb*) must adapt to a changing environment that is shaped by the developing immune response. This necessity to adapt is evident in the flexibility of many aspects of *Mtb* metabolism, including a respiratory chain that consists of two distinct terminal cytochrome oxidase complexes. Under the conditions tested thus far, the *bc*_*1*_*/aa*_*3*_ complex appears to play a dominant role, while the alternative *bd* oxidase is largely redundant. However, presence of two terminal oxidases in this obligate pathogen implies that respiratory requirements might change during infection. We report that the cytochrome *bd* oxidase is specifically required for resisting the adaptive immune response. While the bd oxidase was dispensable for growth in resting macrophages and the establishment of infection in mice, this complex was necessary for optimal fitness after the initiation of adaptive immunity. This requirement was dependent on lymphocyte-derived interferon gamma (IFNγ), but did not involve nitrogen and oxygen radicals that are known to inhibit respiration in other contexts. Instead, we found that *ΔcydA* mutants were hypersusceptible to the low pH encountered in IFNγ-activated macrophages. Unlike wild type *Mtb*, cytochrome *bd*-deficient bacteria were unable to sustain a maximal oxygen consumption rate (OCR) at low pH, indicating that the remaining cytochrome *bc*_*1*_*/aa*_*3*_ complex is preferentially inhibited under acidic conditions. Consistent with this model, the potency of the cytochrome *bc*_*1*_*/aa*_*3*_ inhibitor, Q203, is dramatically enhanced at low pH. This work identifies a critical interaction between host immunity and pathogen respiration that influences both the progression of the infection and the efficacy of potential new TB drugs.

**Author Summary:** Tuberculosis, caused by *Mycobacterium tuberculosis* (*Mtb*) is a serious global health problem that is responsible for over one million deaths annually, more than any other single infectious agent. In the host, *Mtb* can adapt to a wide variety of immunological and environmental pressures which is integral to its success as a pathogen. Accordingly, the respiratory capacity of *Mtb* is flexible. The electron transport chain of *Mtb* has two terminal oxidases, the cytochrome *bc*_*1*_*/aa*_*3*_ super complex and cytochrome *bd*, that contribute to the proton motive force and subsequent production of energy in the form of ATP. The *bc*_*1*_*/aa*_*3*_ super complex is required for optimal growth during infection but the role of cytochrome *bd* is unclear. Here we report that the cytochrome *bd* oxidase is required for resisting the adaptive immune response, in particular, acidification of the phagosome induced by lymphocyte-derived IFNγ. We found that the cytochrome *bd* oxidase is specifically required under acidic conditions, where the *bc*_*1*_*/aa*_*3*_ complex is preferentially inhibited. Additionally, we show that acidic conditions increased the potency of Q203, a cytochrome *bc*_*1*_*/aa*_*3*_ inhibitor and candidate tuberculosis therapy. This work defines a new link between the host immune response and the respiratory requirements of *Mtb* that affects the potency of a potential new therapeutic.

## Introduction

Tuberculosis (TB) is responsible for an estimated 1.4 million deaths annually and remains one of the most deadly infectious diseases (1). The causative agent of TB, *Mycobacterium tuberculosis* (*Mtb*), is an obligate aerobe and relies on oxidative phosphorylation (OXPHOS) via the electron transport chain (ETC) and glycolysis for energy production. The mycobacterial ETC has two terminal oxidases, the cytochrome *bc*_*1*_*/aa*_*3*_ super complex that is related to mitochondrial complex III and IV, and the cytochrome *bd* oxidase which is unique to prokaryotes. These terminal oxidases transfer electrons from the ETC to O_2_ and contribute to the proton motive force (PMF) gradient that powers the production of ATP by ATP synthase. In *Mtb*, cytochrome *bc*_*1*_*/aa*_*3*_ is required for optimal growth and persistence in macrophage and infection mouse models using both genetic (2) and chemical inhibition of the complex (3-5).

In the absence of cytochrome *bc*_*1*_*/aa*_*3*_, electrons are rerouted through the cytochrome *bd* oxidase (6). The latter complex in *Mtb* is encoded in a single operon, *cydABDC*, which produces both the *cydAB* oxidase complex and *cydDC*, a putative ABC-transporter that has not been studied in *Mtb*, but is necessary for assembly of the cytochrome in *Escherichia coli* (7, 8). Genetic deletions in the *cydABDC* operon produce hyper-susceptibility to cytochrome *bc*_*1*_*/aa*_*3*_ inhibitors, demonstrating a partially-redundant role for the terminal oxidases (4, 6, 9). However, the specific role played by the cytochrome *bd* oxidase in *Mtb* remains unclear. In *E*.*coli*, the cytochrome *bd* oxidase detoxifies peroxide radicals and maintains respiration under hypoxic conditions (10, 11). Similarly, studies in the saprophyte, *Mycobacterium smegmatis*, show that *cyd* mutants are hyper-susceptible to peroxide stress and expression of the *cydAB* operon is induced in hypoxic conditions (12, 13). While it is plausible that these properties contribute to *Mtb* fitness during infection, the role played by the cytochrome *bd* oxidase in the mouse model of TB remains unclear. Some studies report no effect of *cydABDC* mutation, while others describe a fitness defect at the later stages of infection (2, 4, 14). Thus, while it is clear that the cytochrome *bd* oxidase is active in mycobacteria, the non-redundant role of this system during infection is unknown.

As an obligate aerobe it is likely that this flexible respiratory chain has evolved to adapt to changing oxygen gradients encountered during infection. During the initial days after infection of the lung, *Mtb* replicates in macrophages, but once these cells are stimulated by T cell-derived cytokines, they restrict *Mtb* growth. Although *Mtb* is exposed to a variety of host pressures during infection, the stressors associated with activation of the macrophage cause a number of specific alterations in the bacterial environment that may alter respiratory requirements. In particular, IFNγ induces antimicrobial responses in the macrophage, including the production of the known respiratory poison, nitric oxide (NO) (15) via nitric oxidase synthase 2 (Nos2). Additionally, inhibition of respiration alters the sensitivity of *Salmonella* to IFNγ-induced superoxide production via the NADPH-dependent phagocyte oxidase (Phox) system (16). Lastly, IFNγ promotes the maturation of the pathogen-containing vacuole, promoting both its acidification and fusion with more degradative compartments. The observation that *cydAB* expression peaks with the onset of the adaptive immune response in the mouse model of infection further suggests an association between T cell cytokines, such as IFNγ, and alterations in the respiratory requirements of *Mtb* (14).

In this work, we investigated the interactions between macrophage activation and the mycobacterial respiratory chain. We report that cytochrome *bd* oxidase is specifically required for the bacillus to resist IFNγ-induced macrophage function. In particular, cytochrome *bd* oxidase is necessary in acidic environments similar to those encountered in the phagosome of IFNγ activated macrophages. These compartments can reach pH levels as low as 4.5 (17, 18), which we show preferentially inhibits the function of cytochrome *bc*_*1*_*/aa*_*3*_ complex. The relative acid-resistance of the cytochrome *bd* oxidase explains its specific function in counteracting IFNγ-dependent immunity.

## Results

### Δ*cydA* mutant is susceptible to IFNγ-activation of macrophages independent of Nos2 and Phox

To investigate the effects of macrophage activation state on the requirement for the cytochrome *bd* oxidase in *Mtb*, we constructed a Δ*cydA* deletion mutant in H37Rv (19). Consistent with previous studies, there was no difference in growth between H37Rv and Δ*cydA* mutants in broth culture (2, 9) (Figure 1A). We compared the fitness of H37Rv and Δ*cydA* mutants in bone marrow-derived macrophages (BMDMs) from C57BL/6J (wildtype) mice. Initially, we used flow cytometry and fluorescent live/dead reporter strains of *Mtb* to estimate relative bacterial growth and viability. The *Mtb* strains expressed a constitutively expressed GFP marker and an anhydrotetracycline (ATc)-inducible RFP marker. GFP intensity was used to estimate total infected cell number and ATc-induced RFP intensity served as a surrogate measure of viability and correlates with CFU (20). By these metrics, the growth and viability of H37Rv and the Δ*cydA* mutant were not appreciably different in unstimulated BMDMs (Figure 1B-C).

**Figure 1:**
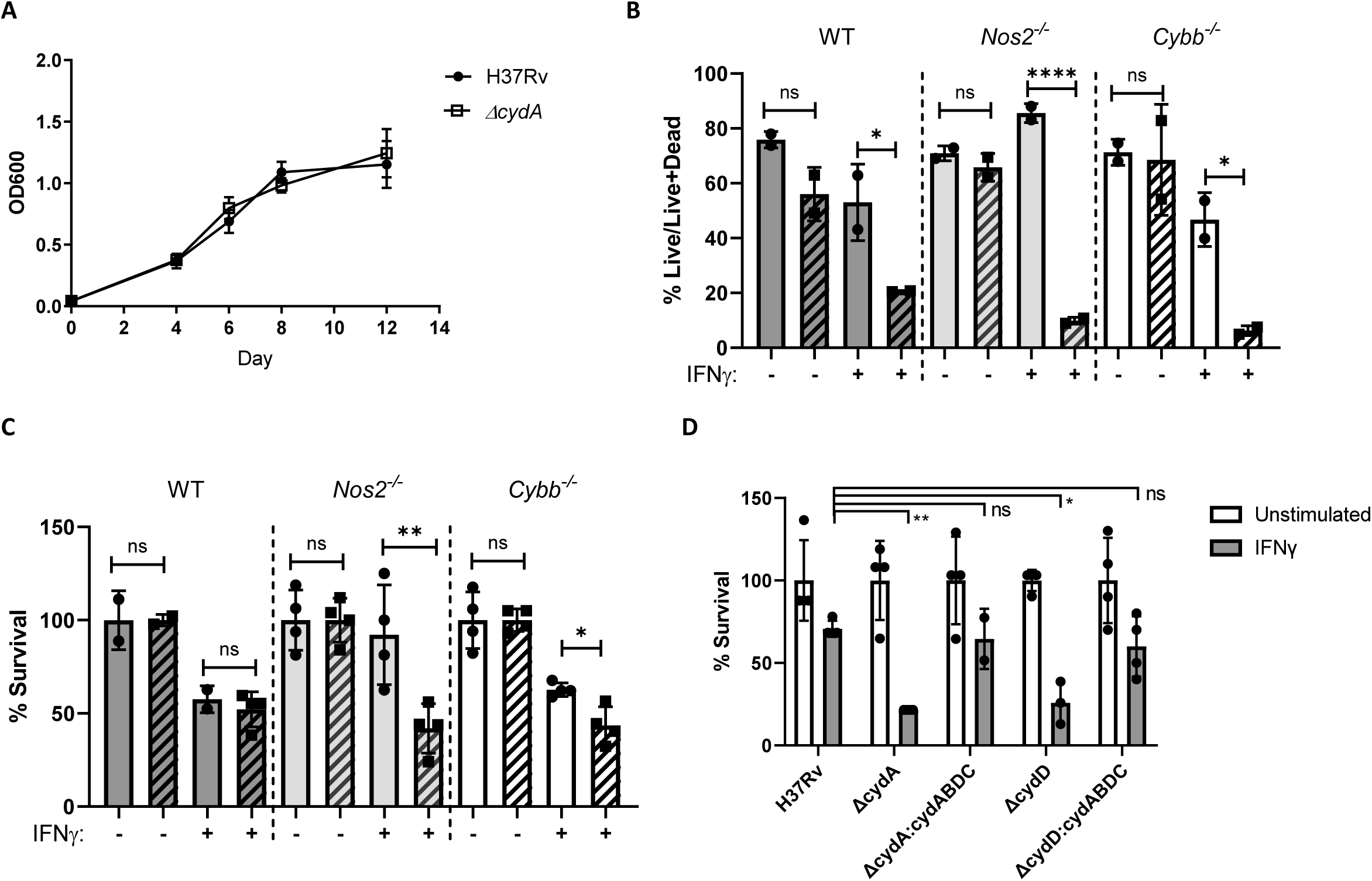
Δ*cydA* mutant is susceptible to IFNγ-activation of macrophages independent of iNOS and Phox. ***cydA* is required to resist IFNγ-mediate immunity independent of Nos2 and Phox.** A) Growth curve of H37Rv and *ΔcydA* grown in broth for 12 days. B) C57BL/6J (WT), Nos2^-/-^ and Cybb^-/-^ BMDMs were left untreated or treated with 25ng/mL of IFNγ for 18 h. Macrophages were infected with H37Rv (solid) or ΔcydA (diagonal bars) live-dead reporter strains (MOI=5). Y-axis represents the fraction BMDMs with live bacteria (RFP^+^GFP^+^) over total infected BMDMs (GFP^+^) determined by flow cytometry. C) IFNγ treated or untreated BMDMs were infected (MOI=5) with H37Rv (solid) or ΔcydA (diagonal bars). CFU determined 4 days post-infection. Represented as percent survival relative to untreated. D) IFNγ treated or untreated Nos2^-/-^ BMDMs infected with H37Rv, Δ*cydA*, Δ*cydD*, Δ*cydA*:Δ*cydABDC*, or Δ*cydD*:Δ*cydABDC* (MOI=5). CFU determined 4 days post-infection. Represented as percent survival relative to untreated. Analysis of B and C was preformed using a two-way ANOVA with Sidak post-test. Analysis of D was performed using a Dunnett’s multiple comparisons test (H37Rv stimulated versus other stimulated conditions). * p-value < 0.05, ** p-value < 0.01, **** p-value <0.0001.

In other bacterial systems, the cytochrome *bd* oxidase is important for resistance to NO and oxidative stress, which are major mediators of IFNγ-dependent antimicrobial activity (10, 21-23). To determine if IFNγ or these reactive species alter the requirement for Δ*cydA*, we stimulated BMDMs with IFNγ, and included cells from *Nos2*^*-/-*^ and *Cybb*^*-/-*^ mice which lack functional Nos2 and Phox systems, respectively. In wildtype BMDMs, addition of IFNγ significantly reduced the number of cells harboring live H37Rv and Δ*cydA* bacteria (Figure 1B). IFNγ treatment had no effect on the viability of H37Rv in *Nos2*^*-/-*^ BMDMs, indicating that IFNγ-mediated inhibition of H37Rv is primarily dependent on NO, consistent with previous studies (18, 22) (Figure 1B). In contrast, *Nos2* and *Cybb* were not necessary for IFNγ to inhibit Δ*cydA* mutants (Figure 1B).

These observations were confirmed by CFU enumeration (Figure 1C). The magnitude of IFNγ-dependent inhibition was smaller in the CFU assay than the flow cytometry study, likely reflecting increased sensitivity of the live/dead reporter for bacterial fitness. Regardless, the CFU assay also showed that IFNγ treatment reduced the viability of H37Rv in a *Nos2*-dependent, *Cybb*-independent manner, whereas the suppression of Δ*cydA* mutants was independent of both mediators. To verify that these mutant phenotypes were due to a lack of *cydABDC* function, we tested a Δ*cydD* mutant and found that it was similarly attenuated in IFNγ-stimulated *Nos2*^-/-^ cells as the Δ*cydA* strain (Figure 1D). Both of these mutants could be complemented *in trans* by expressing the *cydABDC* operon. These observations indicated the cytochrome *bd* oxidase was necessary to resist an IFNγ-induced stress that is independent of Nos2 and Cybb.

### IFNγ but not iNOS or Phox is necessary for the attenuation of *ΔcydA* in mouse lungs

The interaction between IFNγ and Δ*cydA* was then assessed in the mouse model. To evaluate the relative fitness of the Δ*cydA* mutant we performed competition infections using a mixture of H37Rv:Kan and Δ*cydA*:Hyg. To test the importance of lymphocyte-derived IFNγ, the relative fitness of the Δ*cydA* mutant was assessed in wild type C57BL/6J mice and animals lacking adaptive immunity (Rag^-/-^), IFNγ receptor (Ifngr^-/-^), Nos2 (Nos2^-/-^) and Phox (*Cybb*^-/-^) (Figure 2A). At each timepoint, lung homogenates were plated on kanamycin and hygromycin and CFU were enumerated to compare the fitness of H37Rv and Δ*cydA* (Figure 2B). On day 15 post infection, before the onset of adaptive immunity, there was no significant difference in CFU between H37Rv and Δ*cydA* across all five mouse genotypes (Figure 2B). Once the adaptive response was established after 30 days of infection, we observed a significant decrease in Δ*cydA* lung CFU compared to H37Rv in wildtype, *Nos2*^-/-^, and *Cybb* ^-/-^ mice. However, there was no difference between Δ*cydA* and H37Rv lung CFU in *Ifgr*^*-/-*^ and Rag^-/-^ mice demonstrating that the attenuation of the *ΔcydA strain* is dependent on lymphocytes and IFNγ (Figure 2B). Complementation of the *ΔcydA* mutant with *cydABDC* rescued the fitness defect observed in the mutants at the 30-day timepoint in wildtype mice (Figure 2C). These observations were consistent with the *ex vivo* macrophage infections, both indicating that the IFNγ-dependent attenuation of Δ*cydA* is *Nos2* and Phox independent.

**Figure 2:**
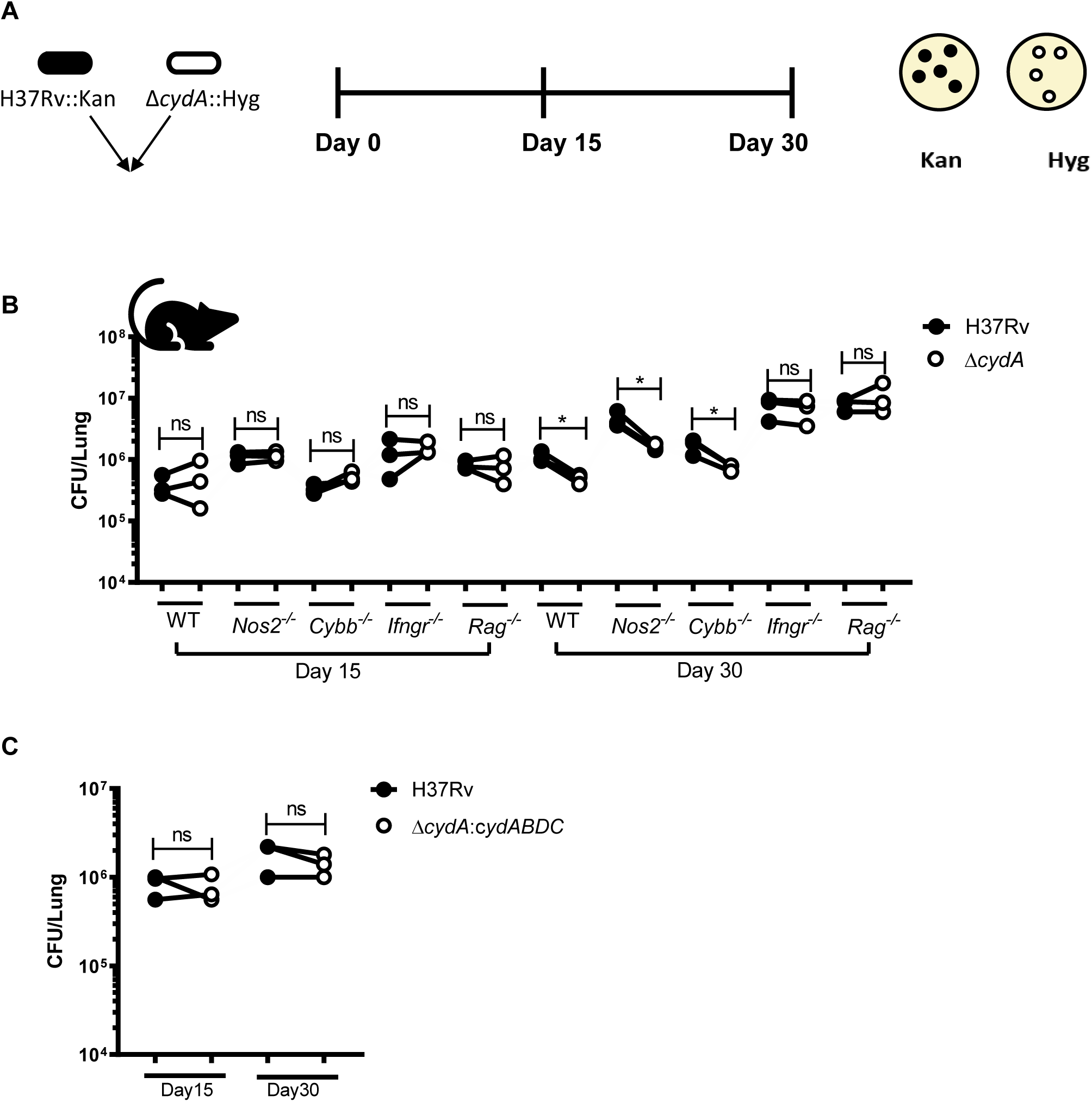
IFNγ but not iNOS or Phox is necessary for attenuation of Δ*cydA* mutants in mouse lungs. ***cydA* is required for persistence in IFNγ competent mice independent of Nos2 or Phox.** A) Schematic of experimental design for 1:1 co-infection with H37Rv::Kan and ΔcydA::Hyg. Lungs were collected on day 15 and day 30 post infection and dilutions of lung homogenates were plated on both 7H10+kanamycin and 7H10+hygromycin. B) Colony forming units (CFU) of H37Rv (black) and ΔcydA (open) in the lungs of C57BL/6J (WT), Nos2-/-, Cybb-/-, Ifngr-/-, and Rag-/- mice were enumerated at day 15 and day 30 post-infection. C) Δ*cydA* mutant can be complemented *in vivo* in C57BL/6J. Lung CFU of H37Rv and ΔcydA:ABDC shown at day 15 and day 30 post-infection.

### *ΔcydA* mutants are defective for growth at low pH

Beyond stimulating the production of RNS and ROS in macrophages, IFNγ also promotes the maturation and acidification of the mycobacterial phagosome. To determine if the hyper-susceptibility of *ΔcydA* to IFNγ could be due to this reduction in pH, we investigated the fitness of *ΔcydA* mutants under acidic conditions. H37Rv and *ΔcydA* were grown in modified 7H9 media at pH values that span those encountered in the maturing phagosome (17, 18). The growth rate of *ΔcydA* was significantly reduced at low pH in comparison to H37Rv or the complemented mutant strain (Figure 3A-C). We next sought to determine if the IFNγ-dependent acidification of the phagosome in macrophages could account for the intracellular growth defect of the Δ*cydA* mutant. BMDMs from wildtype, Nos2^-/-^ and *Cybb*^-/-^ mice were infected with either H37Rv or the Δ*cydA* mutant, and we determined if the inhibitory effect of IFNγ was affected with bafilomycin A1, an inhibitor of the vacuolar-type H+-ATPase that is responsible for the acidification of the phagosome. Using the live/dead reporter assay, we confirmed that Nos2 was required for IFNγ to inhibit H37Rv, but not the Δ*cydA* mutant, providing a situation where the NO-independent inhibitory effect of IFNγ on *bd* oxidase-deficient *Mtb* could be assessed. In these Nos2^-/-^ macrophages, BAF had no effect on the growth of H37Rv, but completely reversed the inhibitory effect of IFNγ on the Δ*cydA* mutant, restoring fitness to levels equivalent to unstimulated BMDMs (Figure 3D). In all three macrophage genotypes, BAF treatment restored the fitness of the Δ*cydA* mutant to wild type levels in the presence of IFNγ. While BAF treatment had a modest effect on the fitness of wild type *Mtb*, the preferential effect on the ΔcydA mutant was consistent with the hypersensitivity of this strain to low pH. Together these observations indicate that the NO-independent inhibitory effect of IFNγ on bd oxidase-deficient *Mtb* could primarily be attributed to the lowered pH of the phagosome.

**Figure 3:**
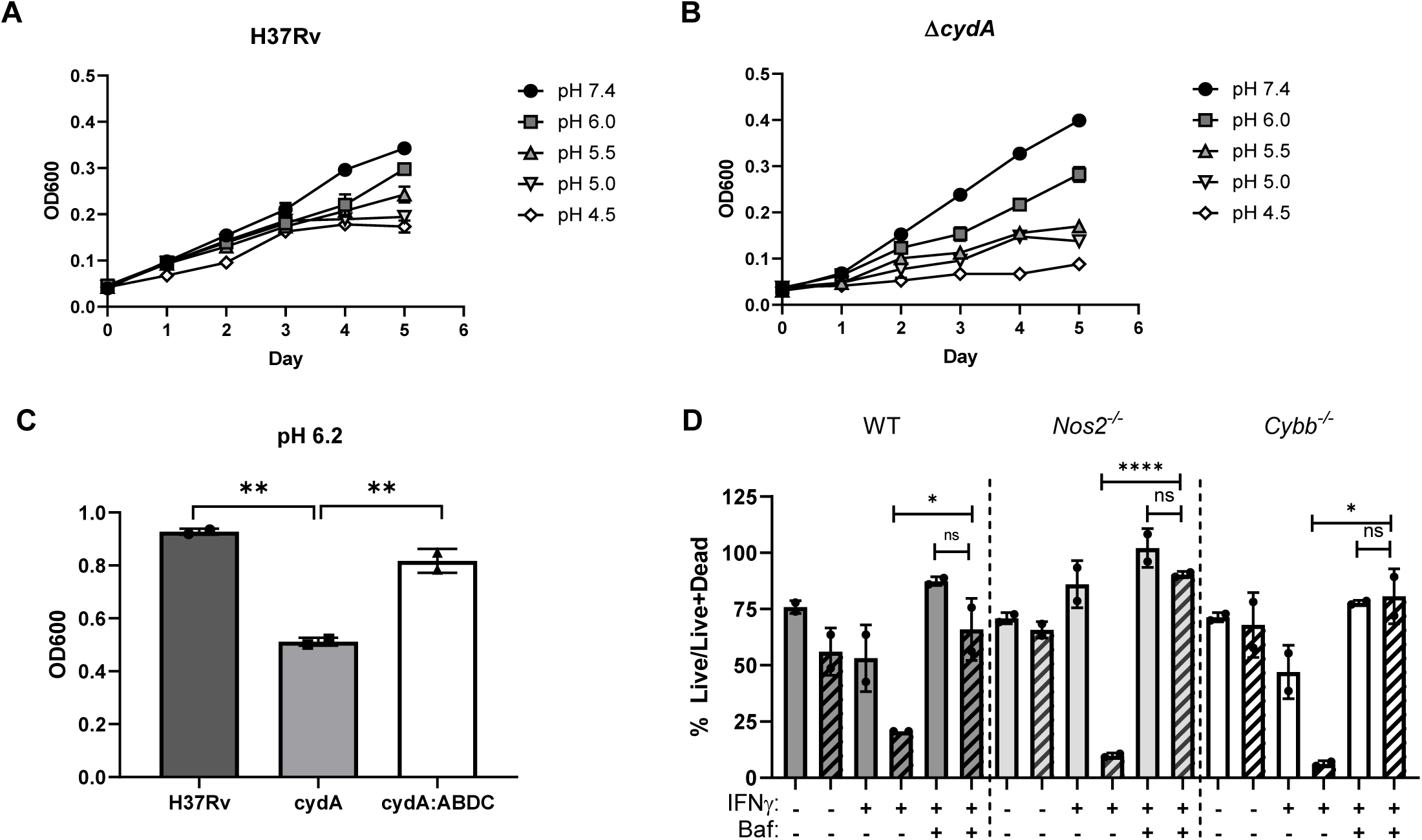
Δ*cydA* are defective for growth at low pH. **cydA is required for growth in acidic conditions.** A, B) Growth of H37Rv (A) and ΔcydA (B) in 7H9-tyloxapol at pH 7.4, pH 6.0, pH 5.5, pH 5.0, and pH 4.5 measured by OD600 for 5 days in a 96 well plate. C) ΔcydA mutant is complemented (ΔcydA:ABDC) at pH 6.2. Cultures grown for 7 days in inkwells. D) WT, Nos2-/- and Cybb-/- BMDMs were left untreated or treated with IFNγ (25ng/mL). Macrophages were infected with H37Rv (solid) or ΔcydA (diagonal bars) live-dead reporter strains (MOI=5). Post-infection BMDMs were left untreated, treated with IFNγ, or treated with IFNγ and Bafilomycin A (100ng/mL). The fraction of macrophages harboring live bacteria (%Live/ Live+Dead) was determined using flow cytometry. Analysis of C was performed using a one-way ANOVA. Analysis of D was performed using a two-way ANOVE with Sidak post-test. Dunnett’s. * p-value < 0.05, ** p-value < 0.01, **** p-value <0.0001

### The *bd* oxidase is necessary for optimal respiration under low pH conditions

Next, we hypothesized that the fitness defect of the Δ*cydA* at low pH was due to reduced activity of the remaining cytochrome *bc*_*1*_*/aa*_*3*_ under these conditions. To evaluate OXPHOS in H37Rv and Δ*cydA* at low pH, we used extracellular flux analysis (Agilent Seahorse XFe96) to measure the oxygen consumption rate (OCR). There was no difference in OCR between H37Rv exposed to pH 7.4 and pH 4.5 media (prior to CCCP addition), suggesting that the respiratory chain in wildtype *Mtb* is able to adapt to reduced pH (Figure 4A). Q203 (Telacebec), an inhibitor of the QcrB subunit of the cytochrome *bc*_*1*_*/aa*_*3*_ complex (5), was used to specifically measure the acid sensitivity of cytochrome *bd* oxidase in H37Rv. At pH 7.4, consistent with previous studies, addition of Q203 led to increased OCR through the re-routing of electrons through cytochrome *bd* (6, 9) (Figure 4A). At pH 4.5, Q203-treated cells displayed an even higher OCR than at neutral pH, potentially indicating further inhibition of the cytochrome *bc*_*1*_*/aa*_*3*_ complex and increased reliance on cytochrome *bd* (Figure 4B).

**Figure 4:**
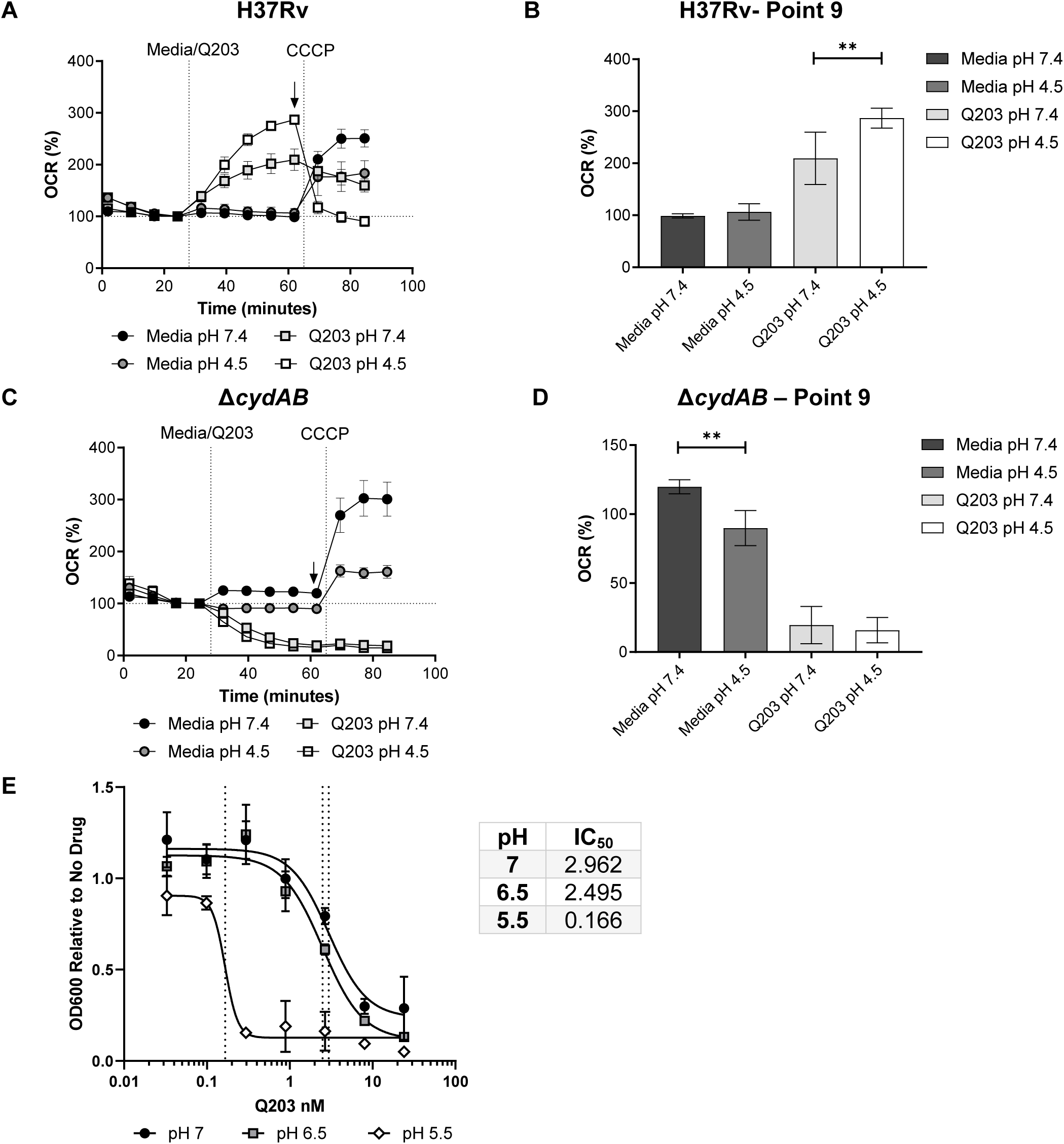
Low pH alters bacterial oxygen consumption rate in Q203 treated WT and *ΔcydAB* strains. **Low pH reduces cytochrome *bc*_*1*_*/aa*_*3*_ dependent oxygen consumption.** A) H37Rv was exposed to media at pH 7.4 and pH 4.5 for ∼30 minutes and then treated with Q203 (10nM), followed by the uncoupler, CCCP. The oxygen consumption rate (OCR) was measured using the extracellular flux analyser B) Bar graphs are plotted from point 9 (indicated by the black arrow in A) before the addition of CCCP. C) Δ*cydAB* mutant was exposed to the same conditions described in A. D) Bar graph of point 9 (black arrow in C). E) IC50 curves of H37Rv treated with Q203 at pH 7.0, pH 6.5 and pH 5.5-day 6. IC50 values for Q203 (nM) at each pH show in the table.

The same experiments were performed with the Δ*cydA* mutant to more specifically probe the effects of low pH on cytochrome *bc*_*1*_*/aa*_*3*_ (Figure 4C). First, we verified that Q203 abrogated OCR in the Δ*cydAB* strain, indicating that both terminal oxidases are inhibited under these conditions. More importantly, decreasing the pH from 7.4 to 4.5 reduced OCR in the Δ*cydA* mutant strain (following media addition – green line and bar), where only the *bc*_*1*_*/aa*_*3*_ complex is active (Figure 4C and D). As this decrease in OCR at low pH was seen only in the Δ*cydAB* mutant and not wild type (Figure 4A), these data indicate that cytochrome *bc*_*1*_*/aa*_*3*_ activity is preferentially reduced under these conditions (Figure 4D). Given this pH-dependent decrease in activity, we hypothesized that acid stress would also increase the sensitivity of cytochrome *bc*_*1*_*/aa*_*3*_ to inhibition. Indeed, reducing the pH from 7.4 to 5.5 enhanced the potency of Q203, lowering the MIC_50_ by almost 20-fold (Figure 4E). In sum, we observed that low pH both decreases *bc*_*1*_*/aa*_*3*_-dependent OCR and increases the potency of a *bc*_*1*_*/aa*_*3*_*-*specific inhibitor. These findings indicate that cytochrome *bc*_*1*_*/aa*_*3*_ activity is inhibited at low pH, and that the cytochrome *bd* oxidase is preferentially required to maintain respiration under acidic conditions, such as those found in the phagosome of IFNγ-stimulated macrophages.

## Discussion

The flexibility of bacterial respiratory chains facilitates adaptation to changing environments, and in many situations the *bd* oxidase becomes critical under conditions where the *bc*_*1*_*/aa*_*3*_ complex is inhibited. In pathogens such as *E*.*coli, Listeria monocytogenes*, and *Salmonella typhimuirium*, the requirement for the cytochrome *bd* oxidase in bacterial virulence has been attributed to its role in resisting hypoxia and nitrosative and oxidative stress (23-26). While previous studies in *M. marinum* and *M. smegmatis* found that the mycobacterial *bd* oxidase can also confer resistance to hypoxia and peroxide, the specific roles played by the cytochrome *bd* and *bc*_*1*_*/aa*_*3*_ oxidases of *Mtb* during infection has been less clear. Our work indicates that the flexibility of the *Mtb* respiratory chain facilitates adaptation to changes in the immune response. As outlined below, our findings specifically suggest that the cytochrome *bd* oxidase provides resistance to IFNγ-mediated immunity by facilitating respiration under the acidic conditions encountered in the phagosomes of IFNγ-stimulated macrophages.

Both in *ex vivo* macrophage cultures and intact animals, we found that the *bd* oxidase was required to resist IFNγ-dependent immunity. These data are consistent with those of *Shi* et al., who showed that *Mtb* cytochrome *bd* oxidase mutants were specifically attenuated in C57BL6 mice only after 50 days of infection. (14). Conversely, other studies have concluded that the cytochrome *bd* oxidase is dispensable for growth in C57BL/6J and BALB/C mice (2, 4). We suspect that these differing conclusions were caused by variations in the infection models. Specifically, our study used a competitive infection model, which ensures that both wild type and mutant bacteria are exposed to identical immune pressures and captures even transient differences in fitness. As a result, we consider competitive studies, such as ours, to be a sensitive approach to detect differences in fitness.

While IFNγ stimulation induces large transcriptional changes and antimycobacterial functions in macrophages, we attributed the *bd* oxidase requirement to alterations in phagosomal pH. Nos2 and Phox are strongly induced by IFNγ, and the requirement for the *bd* oxidase in other bacterial pathogens has been attributed to the resulting NO and ROS (21). As a result, it was somewhat surprising that the sensitivity of Δ*cydA* mutant bacteria to IFNγ treatment was independent of these mediators. Instead, multiple lines of evidence indicated that this mutant was sensitive to the low pH encountered in the phagosome of IFNγ-stimulated macrophages. Firstly, we found that that *bd* oxidase-deficient bacteria grew poorly and respired at reduced rates at low pH. These findings that are consistent with previous transposon mutant screening data suggesting that the *cydABDC* operon was required for optimal growth at pH 4.5 (27). While the reduction in OCR that we observed in the Δ*cydA* mutant at low pH was modest, we speculate that even a small deficit in respiratory rate could cause the observed decrease in growth. These *in vitro* growth defects were related to the intracellular growth environment by demonstrating that inhibition of vacuolar acidification with BAF abrogated the relative fitness difference between Δ*cydA* and wild type *Mtb* in IFNγ-stimulated macrophages. While these data are consistent with a primary role for the *bd* oxidase in adaptation to low pH conditions, we note that inhibition of phagosome acidification can have pleiotropic effects on processes such as phagosome-lysosome fusion and autophagy. Thus, while it remains possible that additional stresses play a role, our observations are consistent with a model in which the *bd* oxidase promotes resistance to the adaptive immune response by promoting respiration in the low pH environment of the IFNγ-stimulated macrophage.

While the requirement for *bd* oxidase activity at low pH can be attributed to the reduced activity we detected for the *bc*_*1*_*/aa*_*3*_ complex under these conditions, the mechanism by which low pH inhibits the activity of the cytochrome *bc*_*1*_*/aa*_*3*_ is unclear. The cytochrome *bc*_*1*_*/aa*_*3*_ super complex is tightly coupled to the transport of protons (28). For every O_2_ molecule reduced by the super complex, 4 protons are pumped into the periplasm and contribute to the proton motive force (PMF) (28). While the cytochrome *bd* oxidase also contributes protons to PMF, it does not have proton pumping capabilities and contributes half of the protons for every molecular oxygen reduced as the super complex (7, 29). It is possible that the tight coupling between proton pumping and electron transfer for the cytochrome *bc*_*1*_*/aa*_*3*_ complex results in its inhibition when extracellular proton concentrations are high. However, acid stress induces a wide variety of transcriptional and physiological responses in *Mtb* and it is also possible that pH has additional indirect effects on the cytochrome *bc*_*1*_*/aa*_*3*_ complex (30-32).

The success of bedaquiline, a mycobacterial ATPase inhibitor, has made respiration an attractive target for new therapeutics. Multiple small molecule inhibitor screens have identified drugs that target the QcrB component of the proton-pumping cytochrome *bc*_*1*_*/aa*_*3*_ (3, 5, 33, 34). Most notably is Q203 (Telacebec) which is currently in clinical trials (5). However, the flexibility of the mycobacterial respiratory chain has raised concerns about the potential efficacy of this drug (2, 4, 35). One strategy to enhance the efficacy of respiratory inhibition is to simultaneously target both the *bc*_*1*_*/aa*_*3*_ and *bd* oxidase complexes, which produces a bactericidal effect (4, 6, 9). Our data suggest that immunity is another important factor that determines the relative importance of terminal oxidases and the ultimate efficacy of these agents. The concept is similar to the previously described synergy between IFNγ-induced tryptophan depletion and the efficacy of *Mtb* tryptophan synthesis inhibitors (27). These examples highlight the importance of understanding the interactions between bacterial physiology and immunity for evaluating and optimizing new therapies.

## Experimental Methods

### Bacterial growth and strain generation

*Mycobacterium tuberculosis* strains were cultured at 37°C in complete Middlebrook 7H9 medium containing oleic acid-albumin-dextrose-catalase (OADC, Becton, Dickinson), 0.2% glycerol, and 0.05% Tween 80 or 0.02% Tyloxapol. Hygromycin, kanamycin, and zeocin were add as necessary at 50 ug/mL, 25 ug/mL, and 25 ug/mL, respectively. All *Mtb* mutant strains were derived from the wildtype H37Rv. Δc*ydA*, Δc*ydD* and Δc*ydABDC* operon were deleted by allelic exchange as described previously (19). The gene deletions were confirmed by PCR verification and sequencing of the 5’ and 3’ recombinant junctions and the absence of an internal fragment within the deleted region. An L5attP-zeoR-cydABDC-operon complementing plasmid was assembled by Gateway reaction (Invitrogen) and transformed into the hygR cydA mutant to generate the cydABDC-complementing strain. The Live/Dead reporter strains were generated by transforming *Mtb* with the replicating Live/Dead plasmid that contains a constitutively expressed GFP and a tetracycline-inducible TagRFP fluorescent protein.

### Ethics Statement and Experimental Animals

C57BL/6 (stock no. 000664), *Cybb*^*-/-*^ (B6.129S-Cybb^tm1Din^/J stock no. 002365), Nos2^-/-^(B6.129P2-Nos2^tm1Lau^/J, stock no. 002609), *Ifngr*^*-/-*^ (B6.129S7-Ifngr1^tm1Agt^/J), and *Rag*^*-/-*^ (B6.129S7-Rag1^tm1Mom^/J, stock no. 002216) were purchased from the Jackson Laboratory. Housing and experimentation were in accordance with the guidelines set forth by the Department of Animal Medicine and University of Massachusetts Medical School Institutional Animal Care and Use Committee (IACUC). Animals used for experimentation were between 6 and 8 weeks old.

### Macrophage infection

Bone marrow derived macrophages (BMDMs) were isolated from C57BL/6, *Cybb*^−/−^ or *Nos2*^−/−^ mice by culturing bone marrow cells in DMEM supplemented with 20% conditioned medium from L929, 10% FBS, 2 mM L-glutamine and 1 mM sodium pyruvate for 7 days. BMDMs were seeded and left unstimulated or stimulated with IFN-γ (25ng/mL, PeproTech) overnight and then infected with *Mtb* at an MOI of 5. After 4 h incubation, macrophages were washed twice with PBS to remove extracellular bacteria and incubated in fresh complete medium with or without IFN-γ. In some conditions, bafilomycin A (100ng/mL, Sigma) or Q203 (at specified concentrations) was added. Cells were lysed with 1% Saponin/PBS (Sigma) at 120 h after infection and then plated on 7H10-OADC plates in serial dilutions. CFUs were counted after 3 weeks of incubation at 37°C.

### Flow Cytometry

For flow cytometry, BMDMs pretreated with or without IFN-γ were infected with Live/Dead reporter *Mtb* strains. At day 3 post-infection, tetracycline (500 ng/ml) was added to medium. Macrophages were harvested after 24 hours tetracycline addition and fixed with 4% PFA for 45 minutes, then run on an LSR II flow cytometer.

### Mouse Infection

Prior to infection, *Mtb* strains were resuspended and sonicated in PBS containing 0.05% Tween80. Δ*cydA* mutant fitness *in vivo* was determined by inoculating mice with a ∼1:1 mixture of Δ*cydA* (hygromycin resistant) and H37Rv (harboring pJEB402 chromosomally integrated plasmid encoding kanamycin resistance) strains via the respiratory route using an aerosol generation device (Glas-Col). At the indicated time points, mice were sacrificed and CFU numbers in lung homogenate were determined by plating on 7H10-OADC agar containing Kanamycin (25 ug/mL) or Hygromycin (50 ug/mL).

### Acid sensitivity assays

The early log-phase of *Mtb* strains were wash twice with PBS and inoculated to a starting optical density at 600 nm (OD_600_) of 0.01 in 96-well plates with 7H9-Tyloxapol-7.4, 7H9-Ty-6.0, 7H9-Ty-5.5, 7H9-Ty-5.0 and 7H9-Ty-4.5, respectively. The desired media pH was achieved by adding 2.5N HCl and 1N NaOH. In some conditions, Q203 (gifted from Professor Barry Clifton) was added. Growth was monitored by OD_600_ daily. Growth rate was determined by comparing OD600 under different pH in day 5.

### Extracellular Flux Analysis

The OCR of Mtb bacilli adhered to the bottom of an XF cell culture microplate (Cell-Tak coated) (Seahorse Biosciences), at 2×106 bacilli per well, were measured using a XF96 Extracellular Flux Analyser (Seahorse Biosciences)(6). All XF assays were carried out in unbuffered 7H9 media (pH 7.4 or pH 4.5 for acidic conditions) without a carbon source. Basal OCR was measured for ∼ 25 min before the addition of compounds through the drug ports of the sensor cartridge. After media or Q203 addition (0.9 µM), OCR was measured for ∼ 40 min, followed by the addition of the uncoupler carbonyl cyanide m-chlorophenyl hydrazone (CCCP) (2 µM) and the OCR measured for a further ∼20 min. All OCR Figures indicate the approximate point of each addition as dotted lines. OCR data points are representative of the average OCR during 4 min of continuous measurement in the transient microchamber, with the error being calculated from the OCR measurements taken from at least three replicate wells by the Wave Desktop 2.2 software (Seahorse Biosciences). The microchamber is automatically re-equilibrated between measurements through the up and down mixing of the probes in the wells of the XF cell culture microplate.

### MIC assay

Log-phase H37Rv was washed twice with PBS + 0.02% tyloxapol and used to inoculate 10mL cultures of 7H9-Ty-7.0, 7H9-Ty-6.5, and 7H9-Ty-5.5 to an OD_600_ of 0.02. As stated before, media pH was achieved by the addition of 2.5N HCl or 1N NaOH. To determine the MIC of Q203 (HY-101040, MedChemExpress), 3-fold serial dilutions from 24nM to 0.3 nM were performed at pH 7.0, 6.5, and 5.5 with a vehicle (DMSO) control. Cultures were incubated at 37°C with shaking. The Syngery HXT microplate reader was used to measure daily OD_600_ of 100uL aliquots in a 96 well plate. The MIC values were calculated on day 6 of growth using nonlinear regression analysis.

## Acknowledgements

We would like acknowledge Kenan Murphy for assisting with mutant generation, and Dirk Schnappinger and Sabine Ehrt for insightful conversations. This work was supported by the National Institutes of Health (grant AI32130 to C.M.S.), and the Arnold and Mabel O. Beckman Foundation (A.J.O.).

